# Evidence of capacitation in the parasitoid wasp, *Nasonia vitripennis* and its potential role in sex allocation

**DOI:** 10.1101/777540

**Authors:** A.R.C. Jones, E.B. Mallon

**Affiliations:** Department of Genetics and Genome Biology, University of Leicester, University Road, Leicester, LE1 7RH, E-mail: /

**Keywords:** sex allocation, sperm motility, transcriptomics, alternative splicing

## Abstract

The allocation of resources to the production of one sex or another has been observed in a large variety of animals. Its theoretical basis allows accurate predictions of offspring sex ratios in many species, but the mechanisms by which sex allocation is controlled are poorly understood. Using previously published data we investigated if alternative splicing, combined with differential expression, were involved with sex allocation in the parasitoid wasp, *Nasonia vitripennis*. We found that sex allocation is not controlled by alternative splicing but changes in gene expression, that were identified to be involved with oviposition, were shown to be similar to those involved in sperm motility, and capacitation. Genes involved in Cholesterol efflux, a key component of capacitation, along with calcium transport, trypsin and MAPKinase activity were regulated in ovipositing wasps. The results show evidence for regulation of sperm motility and of capacitation in an insect which, in the context of the physiology of the *N. vitripennis* spermatheca, could be important for sex allocation.

## Introduction

Understanding the molecular mechanisms controlling an organisms response to their environment, is one of the key questions of biology. A fundamental response to the environment is altering the ratio of male and female offspring, this is, sex allocation (Charnov, 1982; West, 2009). Sex allocation has a large body of theoretical work, and supporting experimental evidence describing its evolutionary role in many taxa (West, 2009). However, outside of circumstances such as temperature dependent sex determination (Göth Ann and Booth David T, 2005; Bull and Vogt, 1979), there is little known about the molecular controls of sex allocation. Frequency dependent selection, was proposed by Fisher to explain sex allocation dynamics (Fisher, 1999). However, when Hamilton derived kin selection, he realised that population level competition in species with limited dispersal, would result in competition between kin. He proposed that selection would act to optimise species sex allocation, in order to minimise competition between kin (Hamilton, 1967). This is Local Mate Competition (LMC) and it provides accurate empirical estimates of optimal sex ratios, based upon population structure. It’s predictions have been supported by observations in mammals, fish and in many invertebrates (West, 2009; Charnov, 1982). LMC is particularly well studied in the parasitoid wasp, *Nasonia vitripennis*, a model for the study of sex allocation.

*Nasonia vitripennis* produces more female biased broods under conditions of high LMC, both in the wild and in laboratory conditions (Werren, 1980, 1983b). Several factors have been found to alter their sex allocation, including the host and brood size (Werren, 1983a; West, 2009). The two main cues females use to alter sex allocation are; 1) if the host has been previously parasitised and 2) the number of local conspecifics (Werren, 1980, 1983a; Shuker *et al.*, 2004). How *N. vitripennis* alter their offspring sex ratio molecularly, has only started to be investigated.

Pannebakker *et al.* 2011 identified three QTL regions linked to offspring sex ratio, along with overlaps with QTLs from clutch size. Three studies have tried to identify gene regulatory changes in sex allocation. Cook *et al.* 2015 and Cook *et al.* 2018 used RNASeq and Pannebakker *et al.* 2013 used a microarray approach, to try and identify key genes that could be involved in sex allocation. The 2015 study used whole bodies and compared three conditions; no host, fresh host and parasitised host to investigate the difference in oviposition and how increased female biased broods in the fresh host compared to the previoously parasitised host. The main gene of interest from the 2015 study and Pannebakker *et al.* 2013 was *glucose dehydrogenase (Gld)*. *Glucose dehydrogenase* is involved in sperm storage in *Drospphilla melanogaster* (Iida and Cavener, 2004). *Gld D. melanogaster* mutants, release sperm at a slower rate then wildtype. With *N. vitripennis* being haplodiploid (fertilised eggs become female, unfertilised eggs become male), the regulation of sperm and fertilisation is key to understanding sex allocation. The 2018 study aimed to identify changes in gene expression in the head, that could be tied to a neurological control of sex allocation using foundress number to alter sex ratios. No differentially expressed genes were identified, indicating that if there are changes in gene expression involved in sex allocation they don’t occur in the brain. The expression of the sex determining splicing factor *double sex* (*dsx*) was altered in relation to oviposition (Cook *et al.*, 2015). Female and male specific splice variants need to be maternally provided for normal sex determination pathways to work (Verhulst *et al.*, 2010, 2013). How this maternal provision is coordinated with sex allocation is unknown.

Alternative splicing describes how mRNA transcripts, from the same gene, can contain different exons and introns resulting in different protein structure and function. This allows for a large variation in protein product to be produced from a single gene. Changing the transcript composition has been shown to be a key regulator in plastic phenotypes, such as between head and body lice, even when no differential gene expression is present (Tovar-Corona *et al.*, 2015).

In the eusocial Hymenoptera, alternative splicing has an important role in reproductive status (Price *et al.*, 2018; Jarosch *et al.*, 2011). We also see alternative splicing involved in neurotransmitter receptors within Hymenoptera (Jin *et al.*, 2007) which have been linked to oviposition in *N. vitripennis* and the regulation of sex allocation by neuronal signalling (Cook *et al.*, 2015). In fact, application of the neurotransmitter acetylcholine agonist imidicloprid to *N vitripennis*, disrupts the detection of the optimal sex allocation (Cook *et al.*, 2016).

We used the 2015 data from Cook *et al.* 2015 and the 2018 data from Cook *et al.* 2018 to investigate alternative splicing and sex allocation. Our aim is to identify if alternative splicing could be involved directly in sex allocation, by searching for an effect caused by foundress number. If we are unable to find an effect caused by foundress number, then we aim to identify any processes that alternative splicing could be involved in regarding either; the regulation and allocation of sperm or epigenetic mechanisms that could be involved in maternal imprinting required for sex determination. We reanalysed the differential expression data using an alignment free approach, as alignment approachs can lead to false positive inflation (Soneson *et al.*, 2016; Bray *et al.*, 2016). We then combined this information from our alternative splicing analysis, using an alignment based approach, to gain as complete a picture of transcriptomic changes in the different treatments as 71 possible.

## Methods

### Structure of the data sets and read processing

The 2018 data set consists of three treatments; single, five and ten foundresses and extracted RNA from the head. The 2015 data is a two by three factorial design with either single or ten foundresses treatments and either no hosts, fresh hosts or previously parasitised hosts. RNA in this study was extracted from whole body samples. SRA files for both of these studies were downloaded from the NCBI database (Accession: GSE105796 and GSE74241) using the e utilities (Sayers, 2017). Reads were then viewed using FASTQC (Andrews, 2014) and then both sets of reads were trimmed using trimmomatic (Bolger *et al.*, 2014) and only the paired reads were used. After looking at the tile quality in the 2015 data set we applied a tile filter using the BBMap functions (Bushnell, 2018).

### Differential expression

Both previous studies had taken an alignment approach to differential expression. We took an alignment free approach using the kallisto 0.43.0 and sleuth 0.30.0 pipeline. We used this approach to reduce the number of false positives (Bray *et al.*, 2016; Pimentel *et al.*, 2017; Soneson *et al.*, 2016). We generated a kallisto index using GCF000002325.3Nvit2rna.fna file from NCBI. The kallisto quantification step was then run with a 100 bootstrap samples. Using the sample run metadata in R (core devlopment team, 2011) and a custom bash script we generated the gene transcript information from the GCF000002325.3Nvit2.1genomic.gtf file. The PCAs for both data sets did not show any outlying samples so none were removed. We then created the full and reduced model and ran the likelihood ratio test, filtering results with a false discovery rate below 0.05. The 2018 data had no significant results with the treatment of foundress number so no further comparisons were made. The 2015 data had no significant results with foundress as the model main effect but when host treatment was used as the models main effect 98 significant results were identified.

### Alternative Splicing

With the *N. vitripennis* genome being sequenced relatively recently and with a high likelihood of new transcripts being identified we chose an alignment approach using the tuxedo suit. We aligned reads to the genome using HISAT2 (Pertea *et al.*, 2016) and sam files were sorted and indexed using samtools (Li *et al.*, 2009). Stringtie was then used to assemble and quantify transcripts, exon and introns using the GCF000002325.3Nvit2.1genomic.gtf file as a guide. A summary using gffcompare found that the 2015 data had 9.4% novel exons and 5.7% novel introns along with 16.9% of loci being novel. Tables of counts were extracted and then read into R (core devlopment team, 2011) and analysed using the ballgown package (Pertea *et al.*, 2016) and visualised using ggplot2 (Wickham, 2009). The PCAs for the 2015 data identified one sample outlier which was then removed from the analysis. Differential transcript usage was determined using FPKM, while differential exon and intron usage was determined using uniquely mapped reads overlapping the exon. Differential transcript usage was identified using the ballgown linear model frame-113 work with a FDR correction, q-values of less then 0.05 were determined as significant.

### Gene ontology (GO) analysis

An annotation of the GCF000002325.3Nvit2 transcriptome was made using trinotate (Bryant *et al.*, 2017). This full GO term list was used to test gene lists of interest for enrichment. Lists of genes were tested for enrichment accounting for GO term structure using the treemap and GSEABase packages (Tennekes, 2017; Morgan *et al.*, 2018) using a hypergeometric test with the GOStats package (Falcon and Gentleman, 2007) and a cut off FDR of 0.05 (Benjamini and Hochberg, 1995).

## Results

### Differential gene expression

With the 2015 and 2018 data, we found that there was no significant clustering for foundress number. For the 2015 data, we did not identify the same strong clustering pattern with host treatment as Cook *et al.* 2015 did.

Neither data set showed any differential expression when foundress was used as the predictor variable. However, as with Cook *et al.* 2015, we identified differential expression with host type as the predictor variable in the 2015 data set which also had host treatment as a variable. We identified 352 differentially expressed genes using host treatment as the predictor variable compared to the 1359 identified in the 2015 data.

### Alternative splicing

No clustering based upon either foundress or host treatment was identified using the PCA of the covariance of the transcript expression in either the 2015 or 2018 data. The 2015 data paper did show some clustering based upon host treatment (figure 1). When using foundress number as the predictor variable to determine differentially expressed transcripts, exons and introns, we only identified one transcript as being differentially expressed in the 2018 data after FDR correction. However no introns or exons from that transcript, or any other, were identified as differentially expressed. We decided this was unlikely to be true differential transcript usage and discarded it. When we used host treatment as a predictor variable, with the 2015 data set, we identified; 1674 transcripts, 3190 introns and 6770 exons that were differentially expressed. We identified 124 genes which had also been identified as differentially expressed from the sleuth pipeline, that also had multiple differentially expressed transcripts. 435 genes were also identified as having differential transcript usage but were not from differentially expressed genes. Differential transcript usage could either be a particular transcript has differntial expression, while all other transcripts in that gene remain the same or it could be isoform switching, where one or several transcripts are upregulated while others are downregulated.

**Figure 1:**
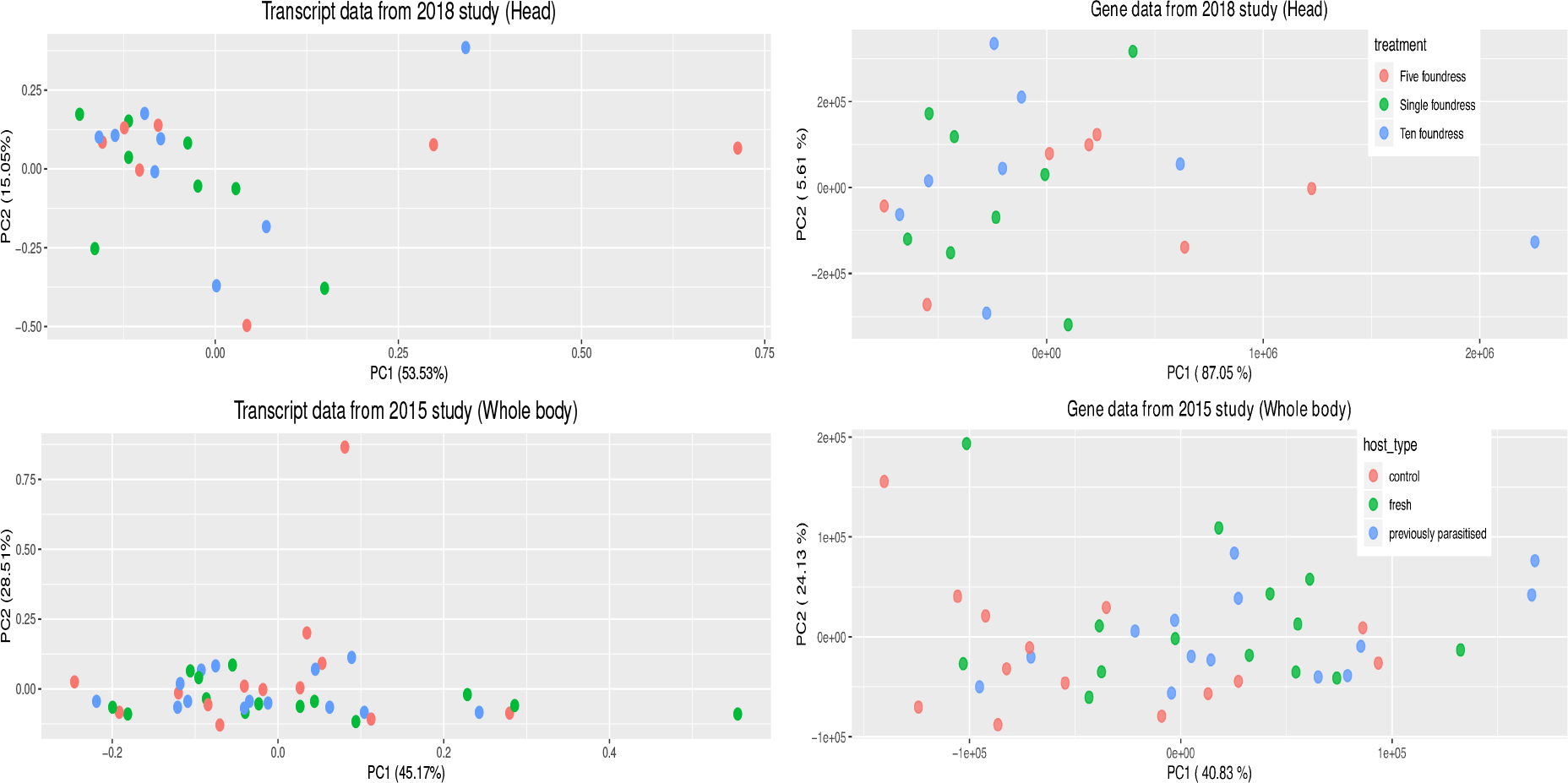
First and second principle components for both (Cook *et al.*, 2018) and (Cook *et al.*, 2015) transcript and gene expression data using FPKM and counts respectively. There is no clear clustering based upon foundress number or host type for either data set.

**Figure 2:**
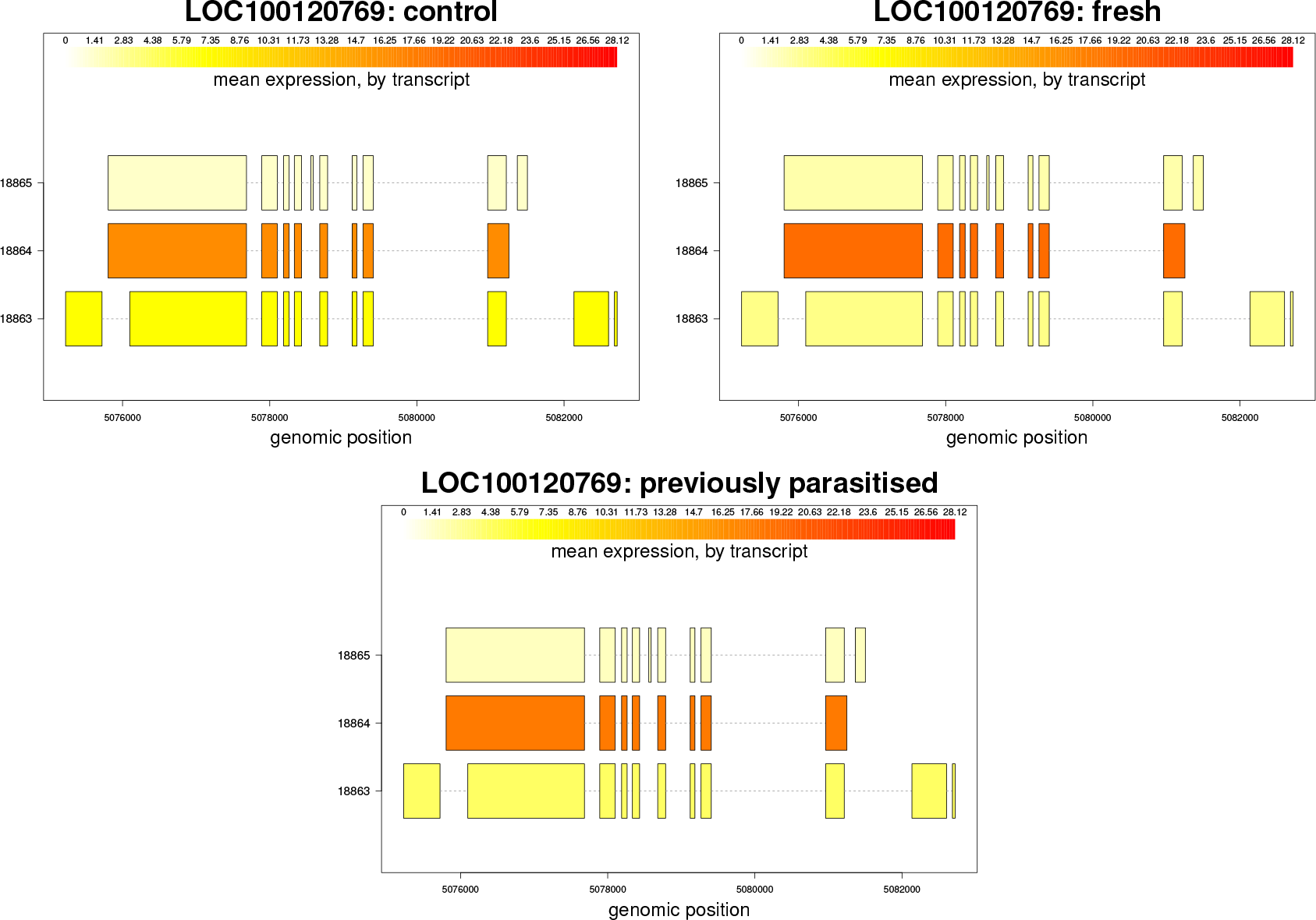
Expression of transcripts in FPKM for *N. vitripennis* LOC100120769 across the three different treatments in the (Cook *et al.*, 2015) data. LOC100120769 is one of the genes with the GO term for a calcium dependent serine/threonine kinase activity.

### GO analysis

To understand the processes being affected by oviposition, we identified enriched GO terms for molecular function, cellular components and biological process for differentially expressed genes, differentially expressed genes which also had differential expression of different transcripts and genes which only had differential transcript usage. For differentially expressed genes we found; 220 enriched GO terms for biological processes, 84 for molecular functions and 38 for cellular components. Within the biological process we see terms for regulation of cardiac muscle contraction, calcium ion regulation and other transmembrance transport. The key terms involved in molecular function include RNA polymerase I initiation, serine protease inhibition and oxidoreductase activity.

174 enriched GO terms for biological processes were identified from genes found to have differential transcript usage as well as being identified as being differentially expressed. This includes terms for positive regulation of insulin receptor signalling pathways, glycoprotein transport, lipid catabolism, negative regulation of prostglandin secretion, mRNA clevage, acyl-CoA biosynthesis and acteyl-CoA metabolism as well as other involved in cell division and organ development. 102 enriched GO terms were identified for molecular function including; methyltransferase activity for histone-glutamine and tRNA, insulin binding and transmembrane transport activity for anions and cholesterol.

For genes that were only identified as having differential transcript usage, we identified 251 GO terms for biological processes, 89 GO terms for molecular function and 40 cellular component terms that were enriched. The biological processes found terms for; regulation of mRNA splicing and processing, snRNA processing, oocyte localisation, regulation of the meitotic cell division and the cell cycle, P granule organisation and germ cell repulsion, negative regulation of histone acetylation and hetrochromatin maintenance involved in silencing. Some of the molecular functions of interest enriched include acetylcholine transmembrane activity, translational repression, RNA binding and MAP kinase activity including calcium modulated MAP kinase activity.

## Discussion

Our findings confirm those found from both Cook *et al.* 2018 and Cook *et al.* 2015, that there is no differential gene expression caused by foundress treatment. We identified fewer genes involved in oviposition than Cook *et al.* 2015, have been able to identify enriched GO terms but did not see the same clustering pattern as Cook *et al.* 2015. This is likely due to the different methods being used with the kallisto sleuth pipeline being more stringent on false positives (Pimentel *et al.*, 2017; Soneson *et al.*, 2016). Our investigation into alternative splicing also identified no significant effect on splicing caused by foundress number. However, by comparing the different gene sets, and those identified in previous studies, patterns that could be informative in determining potential mechanisms regulating sex 185 allocation emerge.

One of the main findings from the Cook *et al.* 2015 was increased expression of *glucose dehydorgenase* (*gld*)(LOC100120817) in ovipositing females. We not only confirmed *gld* is differentially expressed, but also alternatively spliced. *gld* mutant flies were found to be unable to retain the same level of sperm as well as altered the rates of sperm utilisation (Iida and Cavener, 2004). This is not the only gene to be identified that is known to regulate sperm activity. Our study also identified that *Glycerol-3-phosphate dehydrogenase* (GDP, LOC100113822), was alternatively spliced and differntially expressed in ovipositing females. GDP is also involved in calcium dependent lipid metabolism and, in mammals, GDP plays an important part in sperm capacitation, particularly with reactive oxygen species generation (Kota *et al.*, 2009, 2010). The glucose dehydrogenase identified is the FAD-quinone like dehydrogenase, which operates in the absence of oxygen (Tsu-jimura *et al.*, 2006). Another aspect of sperm storage is reducing oxidative stress (Degner and Harrington, 2016), with which we identified several genes including; LOC100123558 an ampdeaminase and LOC103317747 a riboflavin transporter which catalysis oxidation reduction reactions, which were up regulated in ovipositing females. We also identified several other processes which are potentially involved with sperm capacitation.

Capacitation is a series of functional changes which are key for readying sperm for fertilisation. Removing cholesterol from the plasma membrane of sperm, increasing its permeability to bicarbonate and calcium ions, is the defining initial step in capacitation (Ramírez-Reveco *et al.*, 2017). While capacitation has not been identified in insects it has been identified in the mite, *Varroa destructor* (Oliver and Brinton, 1973) and there is some evidence for changes in mosquito sperm (Ndiaye *et al.*, 1997). Two ATP-binding cassette sub-family G member 1-like genes (LOC100123700, LOC100118359) were differentially expressed and spliced while another ATP-binding cassette sub-family G member 1-like gene along with epididymal secretory protein E1-like (a cholesterol transporter) and scavenger receptor class B type 1 (LOC100118508,LOC100115434,LOC100116121) were differentially spliced. These genes are directly involved in cholestrerol efflux. All of these genes were either up regulated in ovipositing females or had specific isoforms that were up regulated in ovipositing females, indicating an increase in cholesterol transport which is synonmous with capacitation.

This change in permeability however requires changes in Ca2+, in order to cause activation in sperm motility. Several enriched GO terms for differential gene expression are involved in calcium and cardiac muscle regulation. There are two sets of genes that seem to be involved in calcium movement. One of these sets were ankyrin homologs, which were down regulated in ovipositing females. These ankyrin genes are involved in regulating Ca2+ in humans, particularly in smooth muscle with a down regulation causing atrial fibrulation (rapid and irregular heartbeats) (Cunha *et al.*, 2011; Le Scouarnec *et al.*, 2008). As these ankyrin genes are predominantly involved in smooth muscle regulation, we believe they are involved in the peristaltic muscle contractions involved in moving developing eggs through the ovariole and oviduct, which have smooth muscle bands (King and Ratcliffe, 1969). Another source of Ca2+could come from the increase in synaptic vesicle glycoprotein 2B-like genes (SV2B, LOC100677929, LOC100122803, LOC100123034, LOC100122992), accompanied by changes in splicing of putative transporter SVOPL (LOC100119949), are more likely to be involved with sperm activation. SV2B is involved in synaptic and neuronal transmission, particulalrly of Ca2+, and studies show that Sv2B knock outs exhibit an elevation of presynaptic Ca2+ levels(Wan *et al.*, 2010; Morgans *et al.*, 2009). As the 2015 data is whole body it is not possible to tell if an increase in calcium is related to sperm activation. The increased SV2B expression is combined with up regulation of EF-hand calcium-binding domain-containing protein 1-like (EFCB1, LOC100117520) in ovispositing females. EFCB1’s cellular component GO term identifies it as been located in the sperm cilia. EFCB1 has been inferred to have an effect on sperm motility and its ortholog has experimental evidence showing an effect on sperm motility in sea squirts (Mizuno *et al.*, 2012).

It is not just the increase in calcium which would indicate that *N. vitripennis* is regulating sperm motility. Sperm have different waveforms; A, B and C which have progressive levels of activity (Thaler *et al.*, 2013). In the mosquito, *Culex quinquefasciatus*, trypsin was identified as inducing the progression to type C motility and is mediated by mitogen activated protein kinase phyosphorylation pathways (Thaler *et al.*, 2013). A similar system has been observed in the common water strider, *Aquarius remigis*, in Lepidoptera, and Orthoptera which would indicate that this is a well conserved system (Miyata *et al.*, 2012; Shepherd, 1974; Aigaki *et al.*, 1987, 1994; Osanai and Baccetti, 1993). Our differential expression analysis, along with Cook *et al.* 2015, identified several trypsin and serine protease inhibitors to be up regulated along with a down regulation of several serine proteases. From our alternative splicing analysis we have identifed 3 trypsin genes that are alternatively spliced (SP97, SP33, SP82) along with a venom serine protease (SP76). Only SP82, a trypsin 1 like endoprotease, was not identified in *N. vitripennis* venom (de Graaf *et al.*, 2010), indicating a role in other biological functions, potently activation of sperm motility.

Our alternative splicing analysis identified several different MAP kinases, including serine/theorine targeting kinase activity, which is important for mediating sperm waveform transition (Thaler *et al.*, 2013), which were calcium/calmodulin dependent. Only one of these MAP kinases had transcripts that were up regulated in ovipositing females, LOC100120769 2. The changing of several MAP kinases splice variants in our data, would indicate a process requiring precise targeting of MAP kinases and the regulation of sperm waveform transitions would fit under that description.

*Nasonia vitripennis* spermathecea consists of an unmuscled capsule that contains sperm, a duct with two bends in it, a muscle that attaches to the duct either side of the bend and a gland with collecting ducts leading into the main sperm duct in the middle of the bend (King and Ratcliffe, 1969). It has been proposed that the bend in the duct acts as a valve, because as the muscles contract, the duct straightens (King and Ratcliffe, 1969) and the duct is small enough to only allow a very limited number of sperm through (Holmes, 1974). The problem with this is the capsule containing the sperm has no musculature to propel the sperm (King and Ratcliffe, 1969). If *N. vitripennis* is indeed able to regulate sperm motility, by capacitation and hyperactivation to waveform C, then the sperm provide the propellant force for the duct to act as a valve. We also see evidence in changes in neurological regulation which could potentially be involved in muscle control.

There is a downregulation of Gamma-aminobutyric acid (GABA) receptor-associated protein, which controls the clustering of GABA receptors. GABA is an inhibitory neurotransmitter and therefore changes in its effect. It would be intriguing to see where this effect is localised, as the spermathecea is located near the terminal ganglion, the largest ganglion in the ventral nerve cord (King and Richards, 2009).

If capacitation is occuring, variation in sensitivity to female controlled sperm activation could also explain the minimal influence of males on fertilisation (Shuker *et al.*, 2006). It could also explain why females with more then one mating have increased first male broods on their first offspring batch, but increased second male broods on the second mating (Boulton *et al.*, 2018). With more sperm available from the second male after the first laying a greater proportion of second male sperm has accesses to the Ca2+ and other enzymes required for activation. What must also be taken into account is that *N. vitripennis* require maternal inputs to successfully develop male or female phenotypes. It would make sense then that these inputs are able to be joined with sex allocation in a complimentary manner.

Maternal imprinting has been identified as being important for *N. vitripennis* sex determination (Verhulst *et al.*, 2010, 2013). In relation to maternally controlled gene expression, our analysis found several genes, identified as being alternatively spliced due to oviposition, involved in RNA processing particularly snRNA, snoRNA and piwiRNA (Werren *et al.*, 2010). In *Caenorhabditis elegans*, recent evidence has shown that snRNA can be transgenerationaly provided, changing gene expression in offspring and these can be neuronally controlled (Ashe *et al.*, 2012; Posner *et al.*, 2019). Our analysis identified alternative splicing of both *Doublesex* and *Transformer2*, which are both maternally provided in *N. vitripennis*. Investigating small RNAs in *N. vitripennis* and identifying if they are maternally provided in a similar manner as seen in *C. elegans* could explain how imprinting and sex allocation could be co-ordinated.

Our main finding is that *Nasonia vitripennis*, during oviposition, displays several changes in gene expression that are known to be involved in regulating sperm motility. These include cholesterol efflux, which is synonymous with mammalian capacitation, as well as changes in trypsin, MAPK activity and calcium regulation. However, as with all whole body studies, there are several different processes occurring that could show similar findings. Further work is needed to understand if there really is a capacitation like process occurring. Given the context, a processes manipulating sperm would be logical. Our findings do offer readily testable predictions which can be experimentally investigated by looking at the location of individual gene expression as well as perturbing expression to see effects on sex allocation.

## Acknowledgements

Big thanks to Cook *et al.* 2015 and Cook *et al.* 2018 for conducting the original studies and generating the great data resource. Thank you to Christian Thomas for reviewing the manuscript. ARCJ was funded by a BBSRC studentship.

## Data Accessibility

The 2015 and 2018 data can be found at NCBI:(Accession:GSE74241,GSE105796) respectively.

## Author contributions

ARCJ and EBM designed the study. ARCJ did the analysis. Both authors wrote and approved the manuscript.

